# Neural correlates of model-based behavior in internet gaming disorder and alcohol use disorder

**DOI:** 10.1101/2023.09.12.557482

**Authors:** Mina Kwon, Hang-Nyoung Choi, Harhim Park, Woo-Young Ahn, Young-Chul Jung

## Abstract

**Background:** An imbalance between model-based and model-free decision-making systems is a common feature in addictive disorders. However, little is known about whether similar decision-making deficits appear in internet gaming disorder (IGD). This study compared neurocognitive features associated with model-based and model-free systems in IGD and alcohol use disorder (AUD).

**Method:** Participants diagnosed with IGD (n=22) and AUD (n=22), and healthy controls (n=30) performed the two-stage task inside the functional magnetic resonance imaging (fMRI) scanner. We used computational modeling and hierarchical Bayesian analysis to provide a mechanistic account of their choice behavior. Then, we performed a model-based fMRI analysis and functional connectivity analysis to identify neural correlates of the decision-making processes in each group.

**Results:** The computational modeling results showed similar levels of model-based behavior in the IGD and AUD groups. However, we observed distinct neural correlates of the model-based reward prediction error (RPE) between the two groups. The IGD group exhibited insula-specific activation associated with model-based RPE, while the AUD group showed prefrontal activation, particularly in the orbitofrontal cortex and superior frontal gyrus.

Furthermore, individuals with IGD demonstrated hyper-connectivity between the insula and brain regions in the salience network in the context of model-based RPE.

**Discussion and Conclusions:** The findings suggest potential differences in the neurobiological mechanisms underlying model-based behavior in IGD and AUD, albeit shared cognitive features observed in computational modeling analysis. As the first neuroimaging study to compare IGD and AUD in terms of the model-based system, this study provides novel insights into distinct decision-making processes in IGD.

## INTRODUCTION

Internet gaming disorder (IGD) encompasses a dysfunctional pattern of gaming characterized by impaired control and elevated priority given to gaming, leading to interference with occupational, social, and academic functions. Previous studies have established similarities between IGD and substance use disorders (SUDs) or gambling disorder in terms of cognitive functioning, neural activities, and other clinical features (for a review, see Vaccaro & Potenza, 2019). Consequently, the World Health Organization officially recognized gaming disorder as a medical condition in the International Classification of Disease-11 (World Health Organization, 2018). However, ongoing debate persists regarding the classification of IGD as an addictive disorder in the same category as SUDs (Kuss, Griffiths, & Pontes, 2017; Saunders et al., 2017; Starcevic, 2017). This debate arises from the distinct clinical aspects of IGD compared to SUDs (Dowling, 2014; Starcevic, 2017), such as less prominent physiological withdrawal and tolerance, which can be attributed to the absence of pharmacological effect in IGD (King, Herd, & Delfabbro, 2017; Yen, Lin, Wu, & Ko, 2022). Therefore, it is critical to identify shared and distinct neurobiological mechanisms of IGD and other substance-related addictive disorders.

According to the reinforcement learning model of addiction, addiction is a transition from goal-directed behaviors into stimulus-driven habitual behaviors (Everitt & Robbins, 2005; Lüscher, Robbins, & Everitt, 2020). Goal-directed behaviors are regulated by the “model-based” system, in which the agent computes and compares possible actions based on the outcomes of each action (Daw, Niv, & Dayan, 2005; Dayan & Niv, 2008). In contrast, habitual behaviors depend on “model-free” system, which relies on previously learned associations between reward and outcome (Daw et al., 2005; Dayan & Niv, 2008; Schultz, Dayan, & Montague, 1997). Thus, model-free decision-making is usually faster and more efficient compared to model-based decision-making but, at the same time, rigid and inflexible. While the model-based and model-free systems cooperatively and competitively arbitrate to make optimal decisions maximizing cumulative rewards (Daw et al., 2005; Drummond & Niv, 2020; Lee, Shimojo, & O’Doherty, 2014), numerous literatures demonstrated impairments in the model-based system and an overreliance on the model-free system in individuals with addiction (Groman, Massi, Mathias, Lee, & Taylor, 2019; Lucantonio, Caprioli, & Schoenbaum, 2014; Voon, Reiter, Sebold, & Groman, 2017). For example, individuals with alcohol use disorder (AUD) and binge drinkers exhibited deficits in model-based control (Chen et al., 2021; Doñamayor, Strelchuk, Baek, Banca, & Voon, 2018; Sebold et al., 2014, 2017; Voon et al., 2015), and severity of alcohol addiction was negatively correlated with model-based behavior in the general population (Gillan, Kosinski, Whelan, Phelps, & Daw, 2016). Furthermore, individuals with gambling disorder, a behavioral addiction without the confound of substance’s neurotoxicity, also showed impaired model-based control (Wyckmans et al., 2019).

While these findings indicate impaired model-based systems as a common psychopathology in addiction, other studies did not report such impairments in individuals with AUD (Voon et al., 2015), high familial AUD risk groups (Reiter, Deserno, Wilbertz, Heinze, & Schlagenhauf, 2016), and young social drinkers (Nebe et al., 2018). Moreover, to the best of our knowledge, the model-based system related to IGD has not been investigated yet. Therefore, to examine whether IGD has similar alterations in model-based systems, it is crucial to compare neurocognitive mechanisms of IGD with those of other substance-related addictive disorders in terms of model-based control. As model-based behavior could be influenced by various factors such as task structure, task instruction, and participants’ cognitive capability (Schad et al., 2014; Silva & Hare, 2020), neuroimaging studies that simultaneously compare different types of addiction are required.

In this study, we investigated model-based and model-free systems in individuals with IGD, individuals with AUD, and healthy controls (HC) using a two-stage task that can assess the balance between the two systems. By applying computational modeling, we aimed to compare latent cognitive processes regarding model-based control in IGD and AUD. Additionally, we used model-based functional magnetic resonance imaging (fMRI) analysis to compare neural activities associated with model-based and model-free systems in IGD and AUD. We hypothesized that (1) model-based behavior would be impaired both in individuals with IGD and AUD compared to HC; and (2) the neural correlates of impaired model-based behavior would differ between IGD and AUD due to the distinct neurobiological effects of alcohol and gaming behavior.

## METHODS

### Participants

Participants were recruited through community and university-based advertisements in Seoul, Korea. A total of 74 participants were included in the analyses, with fMRI analysis conducted on 69 participants after data quality check. Participants were classified by a board-certified psychiatrist into three distinct groups based on DSM-5 criteria for AUD and IGD (HC, *N* = 30; IGD, *N* = 22; AUD, *N* = 22). See **Supplementary materials** for more details.

### Measures

#### Two-stage task

Participants completed the two-stage task (**Figure 1**), which was developed to assess a balance between model-based and model-free behavior (Daw, Gershman, Seymour, Dayan, & Dolan, 2011), inside the fMRI scanner. The task consists of two stages. In the first-stage, participants made a choice between two options, which determined the subsequent second-stage room (blue or yellow). Transitions from the first to the second stage occurred with fixed probabilities: common transitions with a probability of 0.7 and rare transitions with a probability of 0.3. In the second stage, participants made another choice between two options, followed by a feedback stage where monetary rewards were given probabilistically. A hypothetical participant who only uses a model-based strategy (model-based agent) makes choices based on the task structure, specifically the transition probabilities. On the other hand, a participant using a model-free strategy (model-free agent) makes choices solely based on the outcome of the previous trial.

**Figure 1.**
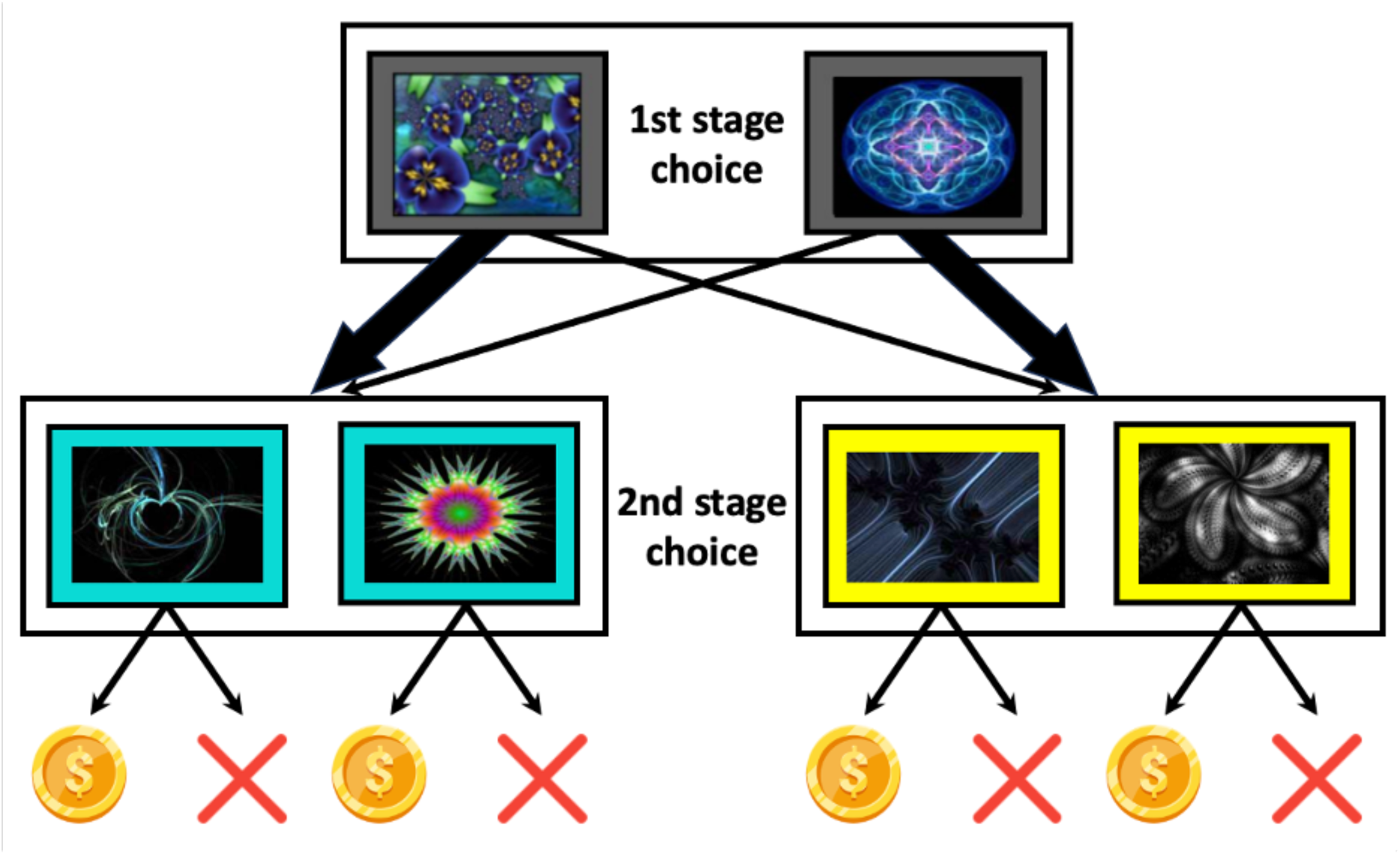
Schematic representation of the Two-stage task. Thick arrows represent common transitions (70%) and thin arrows indicate rare transitions (30%). For example, choosing the left option in the first stage has a 70% chance of transitioning to the blue room in the second stage, and a 30% chance of transitioning to the yellow room. The same probabilities apply in reverse for choosing the right option. After making a choice between the two options in one of the second stages, rewards were given based on slowly varying probabilities ranging from 0.25 to 0.75. Four distinct reward probability distributions were counterbalanced within each group. In the feedback stage, the chosen stimulus from the second stage remained on the screen as a reminder, and outcomes were represented by images of ‘+1000W’ on coins (indicating a reward) or a red ‘X’ (no reward). The value of 1000 won (Korean currency) is approximately 0.76 USD. A hypothetical participant who only uses a model-based strategy (model-based agent) makes choices based on the task structure, specifically the transition probabilities. If a choice in the first stage resulted in a reward through a common transition (70%), the model-based agent repeats the same choice in the next trial. However, if the reward was obtained through a rare transition (30%), the agent switches the choice. On the other hand, a participant using a model-free strategy (model-free agent) makes choices solely based on the outcome of the previous trial; selecting the same option if rewarded and selecting the opposite if not, irrespective of the transition structure.

#### Psychometric measures

Game addiction and alcohol addiction were estimated using modified versions of Young Internet Addiction Test (IAT; Young, 1998) and the Korean version of alcohol use disorder identification test (AUDIT; Saunders, Aasland, Babor, Fuente, & Grant, 1993), respectively. The severity of depressive symptoms and anxiety symptoms were measured using the Beck Depression Inventory (BDI; Beck, Steer, Ball, & Ranieri, 1996) and the Beck Anxiety Inventory (BAI; Beck, Epstein, Brown, & Steer, 1988). For details on other psychometric measures, group differences for each measure (**Table S1**), and correlations between each measure (**Figure S1**), please refer to the **Supplementary materials**.

### Procedure

Participants first completed assessments for the psychometric measures. Then they received instructions for the task and completed the two-stage task inside the fMRI scanner. Lastly, participants underwent an interview with a psychiatrist. Additional details on task instruction, fMRI data acquisition and preprocessing can be found in the **Supplementary materials**.

### Ethics

The study was conducted in accordance with the Institutional Review Board at Severance Hospital, Seoul, Korea (IRB No. 40-2014-0745). All participants were provided with detailed information about the study protocol and provided written informed consent before participating.

### Statistical analysis

#### Behavioral analysis

We conducted a factorial analysis of choice behavior, as replicated from Daw et al. (2011), to calculate stay proportions of first-stage choices at the population level for each group. Linear mixed-effects logistic regression was performed using lme4 package (Bates, Mächler, Bolker, & Walker, 2015) in R to estimate the effects of preceding reward, transition probability, and their interaction on choice behavior. The model included random intercepts and random slopes for the effect of reward and transition probability, with participants as the random factor.

#### Computational modeling

We applied a computational model to the choice behavior based on the hybrid algorithm developed by Gläscher, Daw, Dayan, & O’Doherty (2010), referring to Daw et al. (2011). This model incorporated both model-based reinforcement learning and model-free temporal difference learning. The model-based weight parameter (ɷ) determined the weight given to the model-based learning. Learning rate parameter (α) determines how quickly an agent updates its expected values based on reward prediction errors (RPEs), with higher α indicating faster updates. The perseverance parameter (π) reflects an individual’s tendency to persist with a previously chosen option, regardless of the expected values, with higher π indicating a greater inclination to repeatedly choose the same option consecutively. For each trial, we computed model-free and model-based RPEs, where RPE represents the difference between the actual reward received and the expected reward (Schultz, 2016). We used hierarchical Bayesian analysis for estimating model parameters and group comparisons (Ahn, Krawitz, Kim, Busemeyer, & Brown, 2011; Kruschke, 2014). Additional information regarding the complete equations and details on the computational model and group comparison method can be found in the **Supplementary materials**.

#### Model-based fMRI analysis

In the first-level analysis, we performed a model-based fMRI analysis (O’Doherty, Hampton, & Kim, 2007) to identify voxels of which blood oxygenation level dependent (BOLD) activity is correlated with model-free and model-based RPEs. By using the two parametric regressors of interest, two contrasts were obtained for each participant: one for identifying voxels showing BOLD activity correlated with the model-free regressor, and the other with the model-based regressor. In the first-level analysis, time series of model-free and model-based RPEs were extracted from the computational modeling and included as parametric modulators in the design matrix at the onset of second stage and feedback stage. In the second-level analysis, we conducted two-sample t-tests (HC vs. IGD; IGD vs. AUD; HC vs. AUD) to examine group differences in the neural correlates of model-free and model-based RPE. For more details, please see the **Supplementary materials**.

We then examined correlations between the neural correlates showing significant group differences and the model-based weight parameter (ɷ). This analysis aimed to explore the relationship between model-based behavior and the strength of the neural correlates associated with the model-based and model-free systems, capturing how the strength of each neural correlate varied based on the degree of model-based control. Mean beta values of peak voxels (3mm sphere) in each significant brain region were extracted for each individual, and Pearson correlations were computed with individual estimates of ɷ.

#### Functional connectivity analysis – Psychophysiological interaction analysis

In the model-based fMRI analysis, we found that the insula played a critical role in model-based learning, particularly in the IGD group. To examine how the insula interacts with other brain regions in the context of model-based learning, we conducted a psychophysiological interaction (PPI) analysis (Friston et al., 1997). As insular cortices are known as cortical hubs of the salience network (Seeley, 2019; Seeley et al., 2007), we examined functional connectivity between the bilateral insula and other brain regions related to the salience network (see **Supplementary materials**). In the first-level analysis, the left and right insula (i.e., 3mm sphere centered on peak voxels from the second-level model-based fMRI analysis) were used as seed regions, and the model-free and model-based RPEs were included as parametric modulators. In the second-level analysis, one-sample t-tests were conducted within each group to examine connectivity patterns. We also conducted two sample t-tests (HC vs. IGD; IGD vs. AUD; HC vs. AUD) to examine group differences.

## RESULTS

### Behavioral results

The observed stay probability (**Figure 2**) demonstrates a combination of model-based and model-free learning in all three groups, which is further supported by the results of linear mixed effects logistic regression (**Table S3**). The main effects of reward (HC, *p*<1e-4; IGD, *p*<1e-10; AUD, *p*<1e-06) and transition probability (HC, *p*<1e-9; IGD, *p*<9e-10; AUD, *p*=0.001) were significant in all three groups, indicating that all groups accounted for both rewards (i.e., model-free learning) and the transition structure (i.e., model-based learning) when making decisions. Additionally, the interaction between reward and transition probability also showed a significant effect, supporting the presence of a mixture of model-based and model-free strategies across the groups (HC, *p*<2e-16; IGD, *p*<2e-16; AUD, *p*<2e-16). However, there was no main effect of group in the interaction between reward and transition probability.

**Figure 2.**
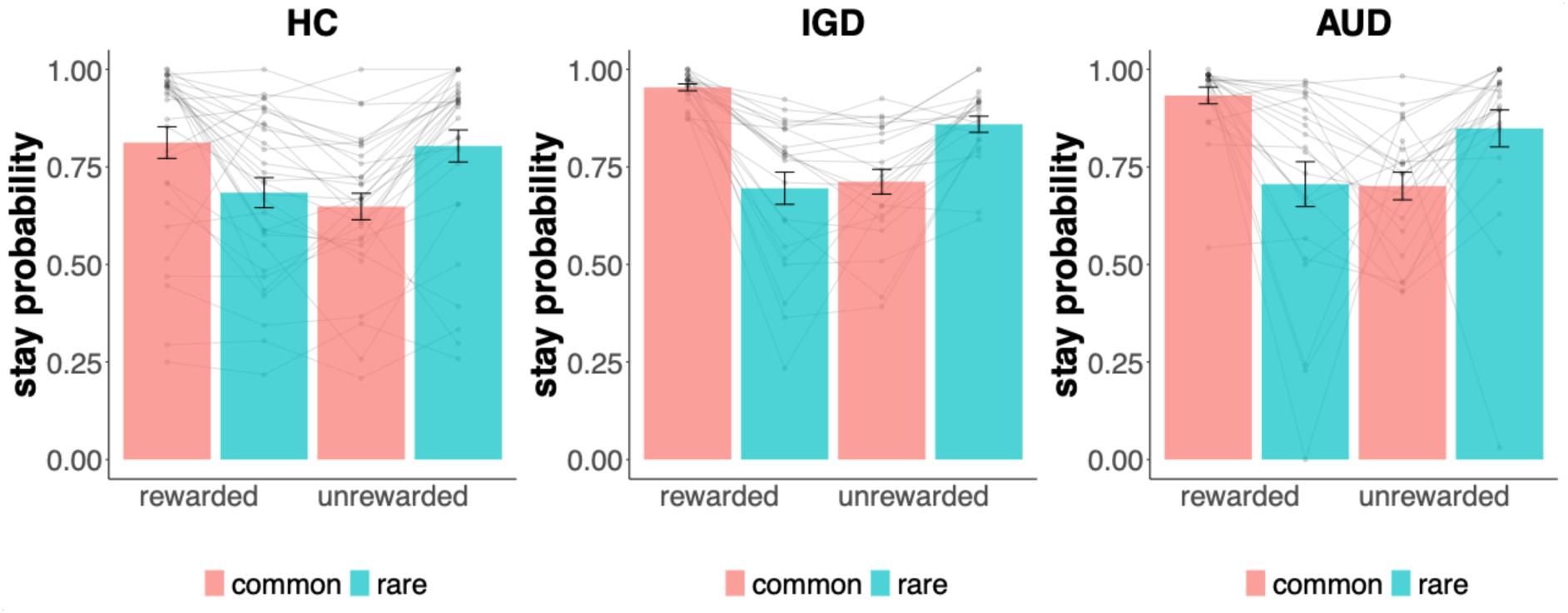
Stay probability of each group. The figure displays the actual stay proportion of each group (HC, healthy control; IGD, internet gaming disorder; AUD, alcohol use disorder) based on reward and transition probability. Each data point represents the individual participant’s data, connected by lines. The error bars represent the standard error of the mean.

### Modeling results

We found significant group differences in the second-stage learning rate parameter *α*_2_, and perseverance parameter π (**Figure 3**). Both the IGD and AUD groups exhibited higher *α*_2_ (*α*_2,*IGD*_− *α*_2,HC_ 95% highest density interval (HDI) = [0.072, 0.359]; *α*_2,*AUD*_ − *α*_2,*HC*_ 95% HDI = [0.011, 0.313]) and higher π estimates (π*_IGD_* – π*_HC_* 95% HDI = [0.100, 1.059]; π*_AUD_* – π*_HC_* 95% HDI = [0.010, 1.021]) compared to the HC group. However, no credible group differences were observed in the other four parameters including the model-based weight parameter ɷ. See **Figure S3** for the distributions of group differences in the model parameter estimates.

**Figure 3.**
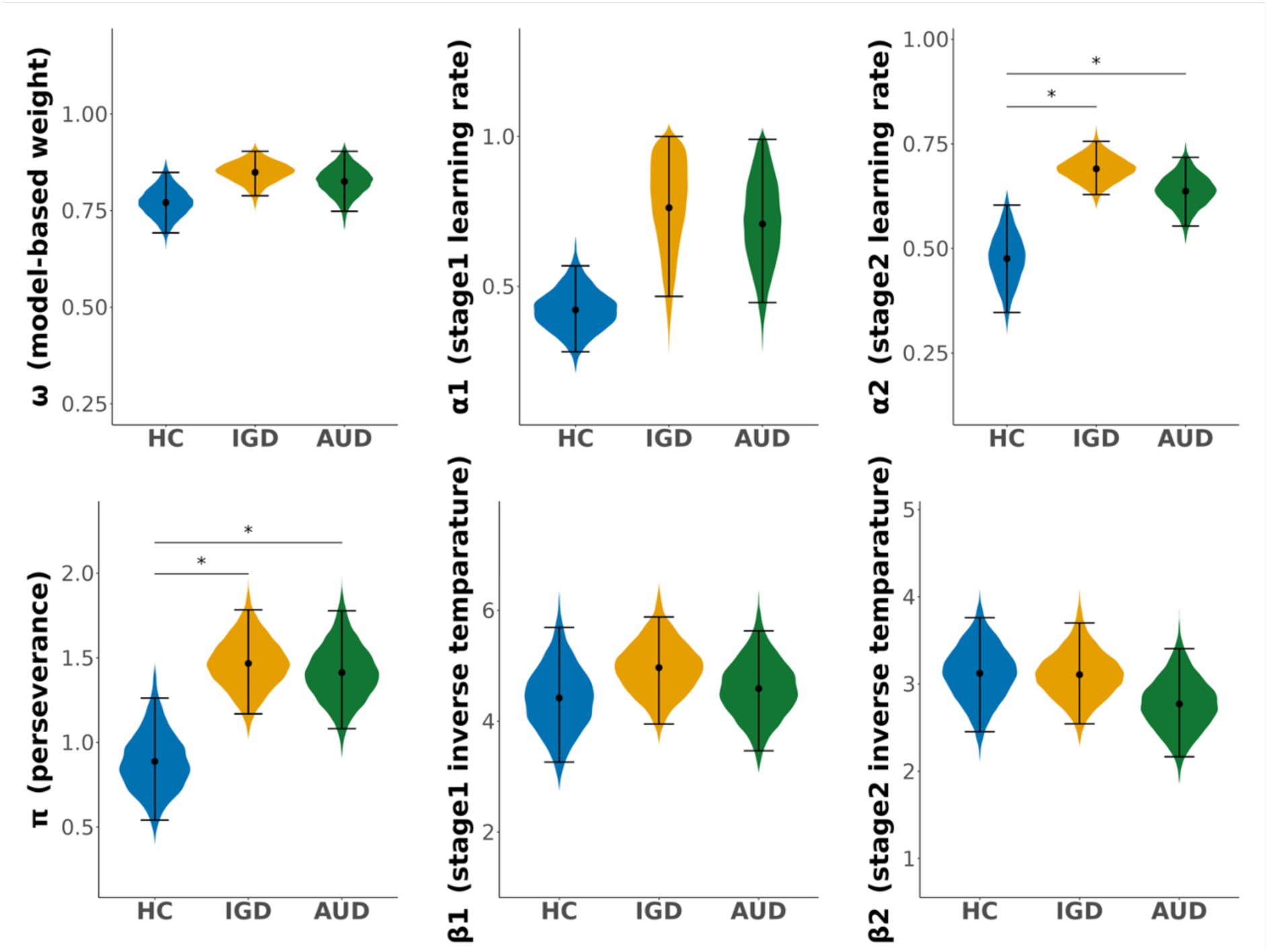
Group comparison of parameter estimates. Each point is a group-wise mean of the estimates of six model parameter (HC, healthy control; IGD, internet gaming disorder; AUD, alcohol use disorder). The model-based weight parameter ɷ represents the degree of model-based control, with higher ɷ values indicating greater emphasis on model-based learning. Learning rate parameters (α_*_, α_!_) indicate how quickly reward prediction errors are updated in the temporal difference learning, independently for stage1 and stage2, with higher values indicating faster updates. The perseverance parameter (π) represents whether the participant has a tendency to choose the same option as on the previous trial, with higher π indicating a greater tendency to repeat choices. Inverse temperature parameters (β_*_, β_!_) determine the level of stochasticity in the participant’s choice, with higher values indicating more deterministic choices. Each error bar represents the 95% highest density interval (HDI) of the parameter estimates. Asterisks indicate significant group differences. See **Figure S2** for the distributions of the group differences.

### Model-based fMRI results

The model-based fMRI analysis revealed significant group difference between the HC, IGD, and AUD groups, when the model-based RPE was employed as a parametric modulator (**Figure 4A & 4B**). Specifically, the activation of the right orbitofrontal cortex (OFC) in the AUD group was greatly correlated with the model-based RPE, compared to the IGD group (*t*=4.22, k=13, *p*<0.001). Furthermore, the bilateral insular activation of the IGD group showed greater correlation with the model-based RPE, compared to the HC group (left: *t*=4.20, k=17, *p*<0.001; right: *t*=4.71, k=33, *p*<0.001). Lastly, activation of the left superior frontal gyrus (SFG) of the AUD group was greatly correlated with the model-based RPE, compared to the HC group (*t*=4.18, k=26, *p*<0.001). However, no significant group differences were observed when utilizing the model-free RPE as a parametric modulator. See **Table S4** for the second level results of each group.

**Figure 4.**
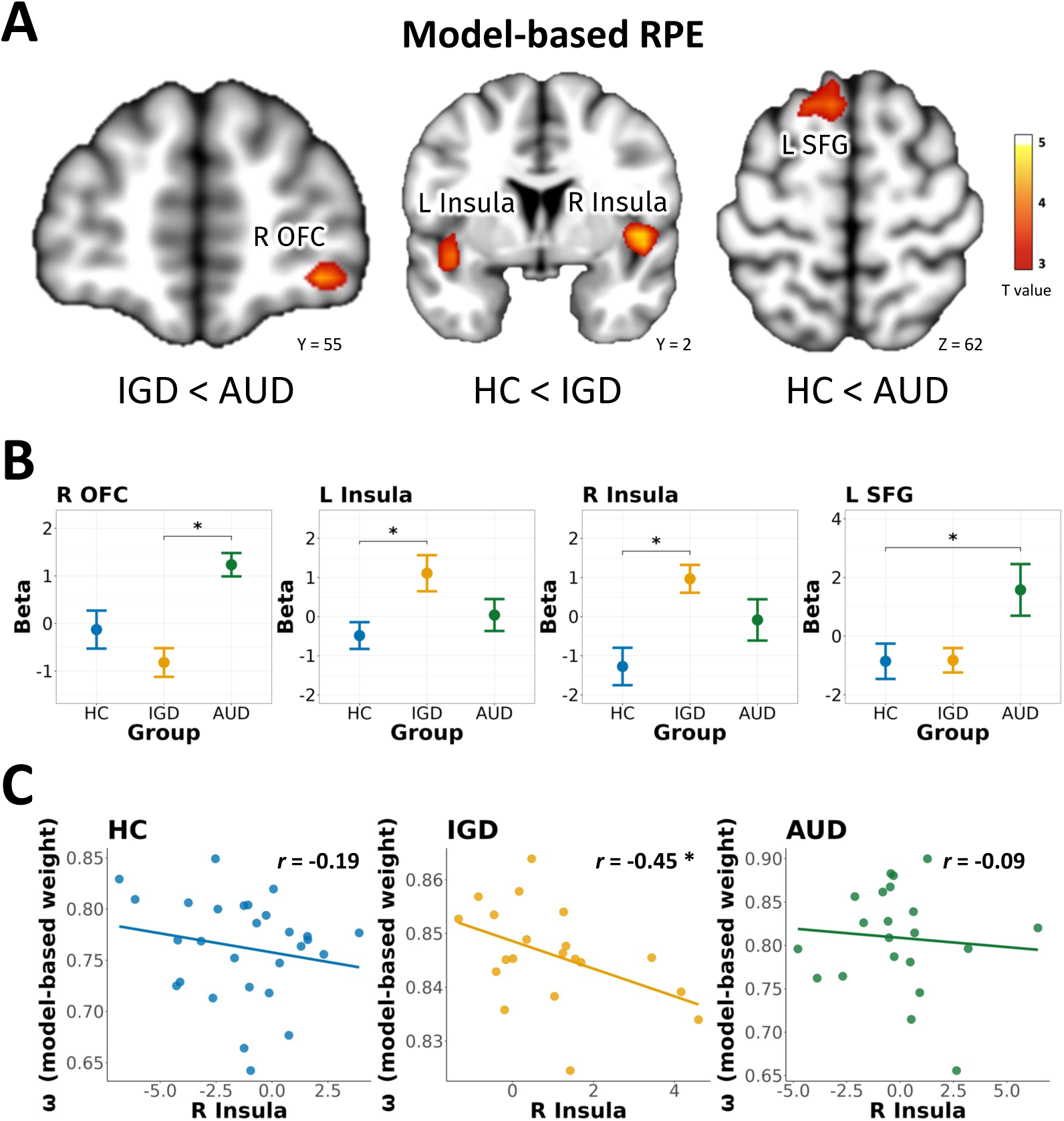
Group differences in model-based fMRI analysis. (A) Brain regions exhibiting significant group differences in the second-level two-sample comparisons using model-based reward prediction error (RPE) as parametric modulators (*p* < 0.001, uncorrected; cluster size, *k* ≥ 10). Significant group differences indicate that one group’s brain activation is significantly correlated with the model-based RPE compared to the other group. The color bar indicates t-statistics from the results of two-sample t-tests. (B) Group-wise average beta value extracted from 3mm spheres at peak MNI coordinates (R OFC: 35, 55, –8; L Insula: –40, 2, –8; R Insula, 46, 2, 2; L SFG: –6, 25, 62) for the brain regions showing significant group differences. Each dot represents the group-wise mean of the beta value for each region, and each error bar indicates the group-wise standard error. Asterisks denote significance based on two-sample t-test (*p* < 0.001). (C) Correlation between beta value of the left insula and the model-based weight parameter (ɷ) estimates. Each dot represents the beta value of the right insula (extracted from 3mm spheres at peak MNI coordinates: 46, 2, 2) from the first-level analysis on the x-axis, and the individual estimates of the model-based weight parameter on the y-axis. The regression line represents the Pearson correlation between the beta value of the right insula and the estimates of ɷ. The asterisk denotes a significant correlation (*p* < 0.05). OFC = orbitofrontal cortex; SFG = superior frontal gyrus.

Further analysis examined correlation between the significant results of the model-based fMRI analysis (i.e., correlation of each region with model-based RPE) and the level of model-based control (e.g., individual estimates of ɷ). We found a negative correlation between the beta value of the right insula and ɷ in the IGD group (*r*=-0.45, *p*<0.05), but not in the AUD group (*r*=-0.09, *p*=0.709) or the HC group (*r*=-0.19, *p*=0.334) (**Figure 4C**). In other words, the coupling between the right insula and the model-based RPE was stronger in individuals with lower model-based weight parameter estimates, only in the IGD group. No significant correlations were found for the left insula.

### PPI results

The PPI analysis revealed significant correlations between the insula and other brain regions in the salience network, only in the IGD group (**Figure 5** & **Table S5**). The right insula was correlated with the right putamen (*t*=5.23, *p*<0.001), left insula (*t*=5.75, *p*<0.001), and occipital lobe (*t*=4.23∼4.37, *p*<0.001) in the context of model-based RPE. Similarly, the left insula was correlated with the anterior cingulate cortex (ACC; *t*=4.24, *p*<0.001), right superior temporal gyrus (STG; *t*=3.67, *p*<0.001), and occipital lobe (*t*=3.72∼5.14, *p*≤0.001). Group comparisons indicated distinctive connectivity patterns in the IGD group (**Table S6**). Connectivity between the right insula and the left insula was greater in the IGD group compared to HC (*t*=3.89, *p*<0.001). Furthermore, connectivity between the insula and occipital lobe was greater in the IGD group compared to the AUD (right insula as seed: *t*=3.52∼4.41, *p*<0.001; left insula as seed: *t=*3.52∼3.57*, p*<0.001) and HC (left insula as seed: *t*=3.41∼3.79, *p*<0.001). Overall, our findings demonstrate that IGD is associated with distinctive patterns of hyper-connectivity in the insula and its interactions with the salience network, suggesting a potential link to ongoing reward processes unique to the IGD group during model-based learning.

**Figure 5.**
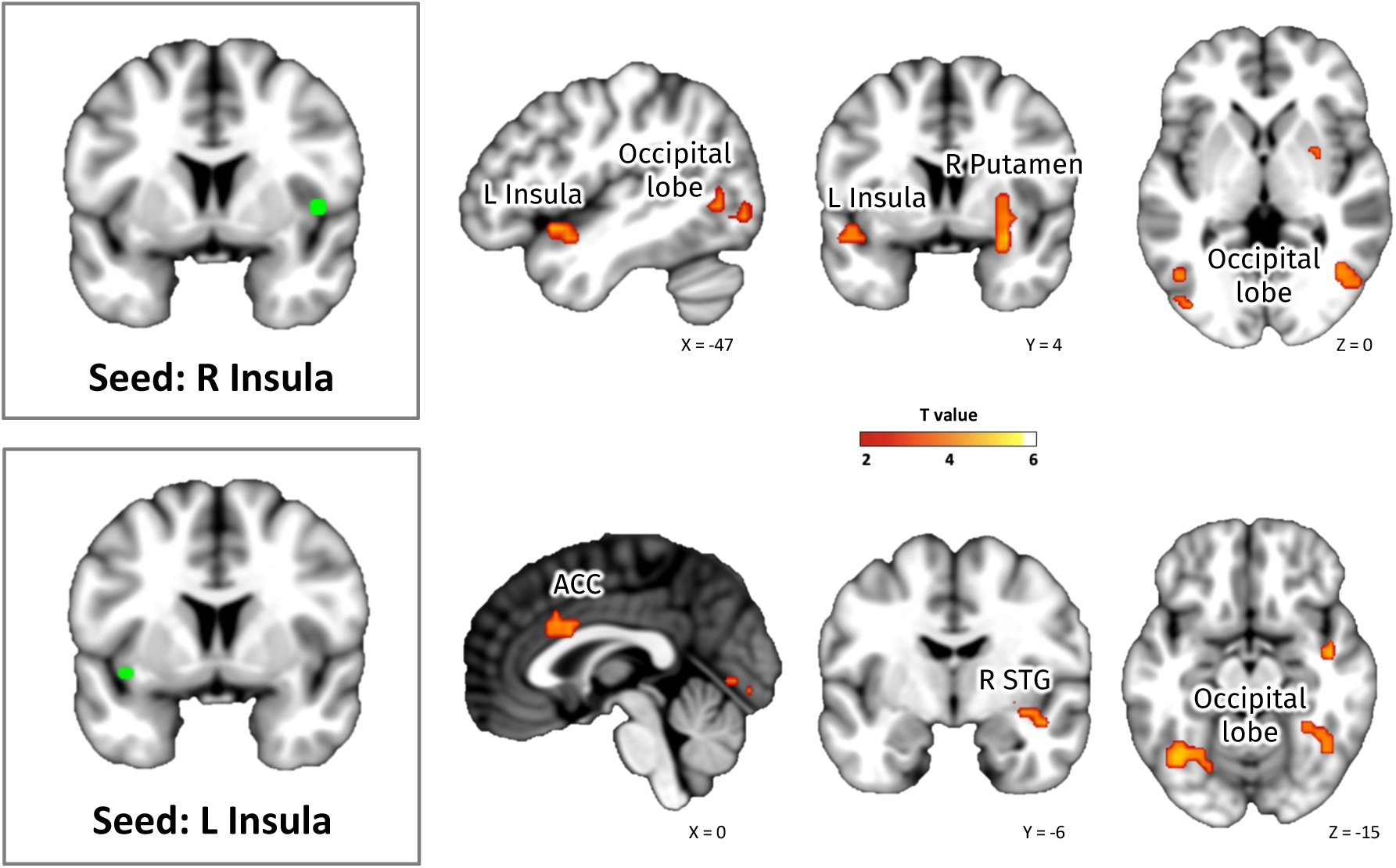
**Results of psychophysiological interaction (PPI) analysis results of the internet gaming disorder group**. Using the right insula (MNI-coordinates: –40, 2, –8) and left insula (MNI-coordinates: 46, 2, 2) as seed regions (3mm spheres), a PPI analysis was conducted with model-based reward prediction error (RPE) as the psychological variable. The brain maps display the effects of model-based RPE on each insula to the whole brain of the IGD group (*p* < 0.001, uncorrected; cluster size, *k* ≥ 10). The color bar represents t-statistics from the results of one-sample t-tests. See **Table S5** for the MNI coordinates of the peak voxels.

## DISCUSSION

In this study, we compared neurocognitive features related to model-based learning in individuals with IGD and AUD. The main findings are as follows: (1) Both the IGD and AUD groups showed higher learning rate and perseverance parameter estimates compared to the HC group, contrary to our initial hypothesis. (2) We found an IGD-specific role of the insula in processing model-based behavior. Model-based RPE was strongly correlated with bilateral insula activation in the IGD group, while it showed correlation with the activation of the frontal regions (i.e., right OFC and left SFG) in the AUD group. Notably, only the IGD group exhibited hyper-connectivity between the bilateral insula and other brain regions in the salience network, including the putamen, ACC, STG, and occipital lobe, in the context of model-based RPE. In addition, we observed that the correlation between the right insula and model-based RPE was particularly stronger in individuals who showed lower model-based behavior (i.e., lower ɷ).

The finding of similarly high model-based behavior across the groups was consistently observed in both behavior and computational modeling analyses. While previous studies have yielded mixed findings regarding the impairment of model-based behavior in individuals with AUD (Chen et al., 2021; Doñamayor et al., 2018; Gillan et al., 2011; Reiter et al., 2016; Sebold et al., 2014; Voon et al., 2015; Wyckmans et al., 2019), a recent study by Silva & Hare (2020) suggested that humans predominantly rely on model-based inference when they have an accurate conception of the task. In our study, we implemented several procedures recommended by Silva & Hare (2020) to ensure a clear understanding of the two-stage task. These procedures included providing a detailed story about the task, administering quiz about the task structure, implementing thorough practice trials, and fixing the stimulus location across trials within each individual. Additionally, participants in our study exhibited above-normal working memory and processing speed, surpassing those reported in previous studies. These higher cognitive capabilities may have contributed to the enhanced model-based behavior observed in our participants (Schad et al., 2014).

While the degree of model-based control (i.e., ɷ) was similar across groups, the IGD and AUD groups exhibited higher *α*_2_ and π estimates, indicating shared cognitive aspects between alcohol and gaming addiction. Impulsivity, a well-known personality trait related to addiction (Bickel & Marsch, 2001; Koob & Volkow, 2016; Kozak et al., 2019; Poulton & Hester, 2019), might account for the higher learning rate (*α*_2_) estimates in terms of reward sensitivity and cognitive control. The faster updates of immediate reward found in the addicted individuals may indicate enhanced salience of immediate rewards or impaired ability to inhibit responses to immediate rewards (Wu et al., 2017). Higher perseverance parameter (π) estimates in the IGD and AUD groups suggest a greater tendency to repeat previous choices regardless of reward, which could be related to habitual behavior or compulsivity, key characteristics of addictive disorders (Everitt & Robbins, 2015; Lucantonio et al., 2014; Lüscher et al., 2020; Ostlund & Balleine, 2009). Therefore, the shared pattern of higher learning rate and perseverance found in the IGD and AUD groups might reflect the neurocognitive characteristics of addictive behaviors.

Using fMRI, we found distinctive neural correlates of model-based behavior in the IGD compared to the AUD groups. Specifically, we observed that the insula plays a specific role in model-based behavior in IGD. The insula is a key component of the brain’s salience network (Menon & Uddin, 2010; Uddin, 2015), involved in representing prediction errors related to reward variance (Preuschoff, Quartz, & Bossaerts, 2008) and processing salient stimuli (Jensen et al., 2007; Seeley et al., 2007). Previous literature reported hyper-connectivity of the salience network in individuals with behavioral addiction during resting state (Tolomeo & Yu, 2022), and also specifically in individuals with IGD during resting state, risky decision making, and executive control tasks (Hong et al., 2015; Lee, Lee, Lee, & Jung, 2017; Sun et al., 2012; Zheng et al., 2019). Consistent with these findings, our results suggest that hyper-sensitivity of the insula, a key component of the salience network, might serve as a distinct neural marker of IGD.

To comprehend the hyperactivation of the insula specifically in the IGD group, it is crucial to consider its involvement in drug craving and addiction (Droutman, Read, & Bechara, 2015). Brain lesions in the insular cortex are known to disrupt addictive behavior, emphasizing the insula’s significant role in addiction (Naqvi, Gaznick, Tranel, & Bechara, 2014). However, neuroimaging studies on SUDs have generally reported hypoactivation of insula during decision-making tasks (Nestor, Hester, & Garavan, 2010; Stewart, Connolly, et al., 2014; Stewart, May, et al., 2014). This discrepancy may be attributed to the differential processing of drug rewards and non-drug rewards in individuals with addiction (Madden, Petry, Badger, & Bickel, 1997). Drugs of addiction are highly salient reward, leading to hyperactivation of the salience network, while non-drug rewards are associated with decreased activity in the salience network compared to non-drug users (Cushnie, Tang, & Heilbronner, 2023). Therefore, the previous findings of hypoactivation of the insula during decision-making tasks in SUDs may be due to the use of non-salient rewards. Correspondingly, neuroimaging studies of gambling disorders using simulated gambling tasks have indicated heightened activation of the salience network (Clark, Lawrence, Astley-Jones, & Gray, 2009; Clark, Studer, Bruss, Tranel, & Bechara, 2014). Considering the task dynamics and reward structure in our study, which may resemble gaming behavior at least more than substance use, the excessive insula activation observed in the IGD group may reflect that the individuals treated the two-stage task like a game.

Another notable finding is that individuals with AUD exhibited hyperactivation in the prefrontal regions, specifically the OFC and SFG, in the context of model-based behavior. The OFC has been identified as a key regulator of goal-directed, or model-based action planning and execution (Gremel & Costa, 2013; Jones et al., 2012; Kahnt, 2023; McDannald, Lucantonio, Burke, Niv, & Schoenbaum, 2011), encoding the value of stimuli and the relationship between stimuli and their expected outcomes (Howard, Gottfried, Tobler, & Kahnt, 2015; Lopatina et al., 2015). Similarly, the SFG is involved in higher cognitive functions and working memory associated with executive processing (Boisgueheneuc et al., 2006). Interestingly, while previous literature suggested decreased prefrontal activation in individuals with AUD during decision-making tasks and resting state, the individuals with AUD in our study showed increased activation of these regions. This discrepancy could be explained by differences in model-based performance, as we did not detect impairment of model-based control in AUD, contrary to previous findings of AUD or SUDs (Dom, Sabbe, Hulstijn, & Brink, 2005; Moorman, 2018). This could be attributed to the task environment that facilitated model-based behavior (Silva & Hare, 2020) and the higher cognitive abilities of the participants in our study (Schad et al., 2014). In summary, the hyperactivation observed in the frontal regions in individuals with AUD may reflect compensatory efforts to counteract impaired cognitive control and decision-making abilities. It is important to note that compensatory hyperactivation in the frontal regions is specific to AUD and not observed in IGD. This distinction may be attributed to the absence of chemical intoxication in IGD, unlike in AUD and other SUDs (Grant, Potenza, Weinstein, & Gorelick, 2010; Han et al., 2015; Oscar-Berman & Marinković, 2007; Weinstein & Lejoyeux, 2020).

To the best of our knowledge, this is the first neuroimaging study that compared IGD and AUD focusing on the model-based behavior. Our findings suggest that IGD and AUD have shared mechanisms regarding the model-based behavior, indicating common cognitive characteristic related to reward sensitivity and compulsivity. However, the underlying neural mechanisms may differ due to various factors such as differences in reward of salience and the distinct brain changes related to alcohol and gaming behavior. Our findings provide valuable insights into the neurocognitive mechanisms underlying addictive disorders and highlight the need for further research to explore the complex role of insula and salience network in IGD. Nonetheless, this study has several limitations. First, the relatively modest sample sizes of each group curtail statistical power, thereby accentuating the importance of replicating these results in future investigations with larger sample sizes and more stringent statistical thresholds. Second, although participants were diagnosed with IGD and AUD by psychiatrists, they were not clinical patients seeking for treatment. Instead, they were recruited from community sites, which could potentially lead to variations in clinical characteristics compared to patients with more severe symptoms. Third, the generalizability of the findings is limited due to the restriction to young men in terms of gender and age. Lastly, the majority of individuals in the IGD group was predominantly engaged in a specific game genre, multiplayer online battle arena (MOBA), which may be influenced by Korean culture. Therefore, future studies considering the influence of culture are warranted.

## CONCLUSIONS

Our study provides novel insights into the neurocognitive features and neural correlates of model-based learning in individuals with IGD and AUD. Despite both IGD and AUD exhibiting similar levels of model-based behavior, distinct neural signatures were observed in the insula for IGD and prefrontal regions for AUD. These findings suggest potential differences in the neurobiological mechanisms underlying addictive behaviors in IGD and AUD, contributing to the growing body of evidence highlighting shared and distinct features of IGD and substance-related addictive disorders. Further prospective research is needed to better understand pathophysiology of IGD, and to address the controversy surrounding the diagnostic criteria and treatment approaches specific to IGD.

