## Supplementary materials for "Neural correlates of model-based behavior in internet gaming disorder and alcohol use disorder"

*Participants*

Participants were recruited from September 2018 to August 2019 through community and university-based advertisements in Seoul, Korea. Initially, 77 participants were enrolled, but three of them were subsequently excluded: one participant did not complete data collection, one participant was excluded due to moderate depression and anxiety disorder diagnosed by psychiatrists, and one participant was excluded due to the presence of comorbidity between alcohol use disorder (AUD) and internet gaming disorder (IGD). Consequently, a final sample of 74 participants were included in the analyses, with functional magnetic resonance imaging (fMRI) analysis conducted on 69 participants after data quality check (see ***fMRI data acquisition and preprocessing** below*). Participants underwent evaluation by a board-certified psychiatrist using a semi-structured interview based on the Structured Clinical Interview from the DSM-IV (SCID; First, Spitzer, Gibbon, & Williams, 1997) to assess major psychiatric disorders. Additionally, the Korean version of the Wechsler Adult Intelligence Scale (K-WAIS IV; Wechsler, 2008) was administered, and individuals with a verbal IQ or performance IQ of less than 80 were excluded. Two psychiatrists classified participants into three distinct groups based on DSM-5 criteria for AUD and IGD (HC, $N$ = 30; IGD, $N$ = 22; AUD, $N$ = 22).

*Psychometric measures*

In addition to the self-report surveys conducted to measure game addiction, alcohol addiction, depressive, and anxiety symptoms, we also assessed impulsivity, childhood attention-deficit hyperactivity disorder (ADHD) symptoms, and intelligence to account for potential group differences. Self-reported impulsivity was evaluated using the Barratt Impulsiveness Scale (BIS; Patton, Stanford, & Barratt, 1995), and childhood ADHD symptoms were assessed using the Wender Utah Rating Scale (WURS; Ward, Wender, & Reimherr, 1993). Verbal IQ and performance IQ were measured using the Korean version of the Wechsler Adult Intelligence Scale (K-WAIS IV; Wechsler, 2008).

To examine group differences in survey measures, we performed a one-way analysis of variance (ANOVA) with group factors (i.e., HC, IGD, AUD) (**Table S1**) and performed post-hoc pairwise comparisons using Tukey's test (**Figure S1**). The IGD group exhibited the highest internet addiction symptoms, while the AUD group showed the highest AUD symptoms. We also found significant group differences in depression, anxiety, impulsivity scores, and childhood ADHD symptoms. Specifically, the IGD group reported higher depression symptoms compared to the other two groups and higher anxiety symptoms compared to the HC group. Impulsivity was highest in the IGD group, followed by the AUD group and the HC group. Additionally, the IGD group reported the highest childhood ADHD symptoms, with a significant difference from the HC and AUD groups. IQ scores, including all subscales, and age did not show significant statistical differences.

*Task instructions*

Participants were instructed that they will be playing a treasure hunt game repeatedly, where earning more treasures would result in higher payment. Based on the schematic figure depicting the task (**Figure S2A**), the instruction stated, "You choose a door from two rooms to find the treasure. First of all, it starts in a gray room with two doors, each distinguished by a unique picture. Choosing the left door has a higher probability of transitioning to the blue room and a lower probability of transitioning to the yellow room, while choosing the right door has a higher probability of transitioning to the yellow room and a lower probability of transitioning to the blue room. Throughout the game, these probabilities are fixed. Here is an example. Let's say you choose the left door in the first room (the gray room), and now you are in the blue room. In the second room, there are two doors, each with distinguishing pictures. Let's say you choose the right door this time. Opening the door will either reveal a "treasure" or result in a miss. The chances of finding the treasure gradually change from 25% to 75% over time, but the specific probabilities are not revealed. Please choose doors carefully to maximize the chances of discovering treasures." Subsequently, based on **Figure S2B**, we explained the entire process of the task once again, including the duration of each stage: 2.5 seconds for choosing the first door, a 1-second wait to move to the next door, 2 seconds for choosing the second door, a 1.5-second feedback check, and a 0.5-second wait for the next treasure hunt.

After the instruction, participants underwent extensive practice trials consisting of 30 trials of first-stage choice followed by transition to the second stage, 10 trials of the second-stage choice followed by the feedback phase, and finally, 67 trials of the complete trial (first- and second-stage choices in a row). Different fractal images were used in the practice trials compared to the actual task. Subsequently, participants were asked to answer three questions to assess their understanding of the task: "The first door is selected in the gray room (O/X)", "When you choose a door in the first room, the probability of transition (to the blue room or yellow room) does not change (O/X)", and "The door with the highest probability of receiving the treasure in the second room changes over time (O/X)."

*fMRI data acquisition and preprocessing*

MRI scanning was performed using a 3.0T MRI scanner (MagnetomVerio, Siemens Medical Solutions, Erlangen, Germany). Functional images were acquired with three 8.5-minutes runs using a T2*-weighted gradient echo-planar imaging sequence (30 axial slices, 4 mm thickness with 1 mm interslice gap; repetition time = 2,000 ms, echo time = 30 ms; flip angle = 90°; in-plane matrix size = 64 × 64 pixels; and field of view = 240 mm) while participants performed the two-stage task. Additionally, a structural T1-weighted gradient echo image was acquired (matrix size = 256 × 256, number of slices = 176, slice thickness = 1 mm, echo time = 2.46 ms, repetition time = 1,900 ms, field of view = 250 mm, flip angle = 9°, bandwidth = 170Hz/Px). Preprocessing of the fMRI data was conducted using fMRIPrep version 20.2.1. This included spatial normalization, susceptibility distortion correction, co-registration, and slice timing correction using fMRIPrep (Esteban et al., 2019). For data quality, participants with an average framewise displacement value higher than 0.5 (Power, Barnes, Snyder, Schlaggar, & Petersen, 2012) were excluded from the analysis ($N$ = 4). Additionally, one participant was excluded from the analysis due to signal loss in the frontal regions. This resulted in a total of 69 participants included in the fMRI analysis (IGD, *N*=20; AUD, *N*=21; HC, *N*=29).

*Computational modeling*

To select the best fitting model, we tested three different models: a seven-parameter model used in Daw, Gershman, Seymour, Dayan, & Dolan (2011), a six-parameter model (see below), and a four-parameter model (Wunderlich, Smittenaar, & Dolan, 2012). After fitting each model separately for each group using hierarchical Bayesian modeling (Rouder & Lu, 2005), we computed the leave-one-out information criterion (LOOIC) using "loo" package in R (Vehtari, Gelman, & Gabry, 2017). **Table S2** presents the LOOIC of each model for each group. As a lower LOOIC indicates a better model fit, the six-parameter model was the best fitting model for all three groups. Consequently, we used the six-parameter model for further analyses.

We fit each participant's trial-by-trial responses in the task with a hybrid reinforcement learning (RL) model described in previous literature (Daw et al., 2011; Gläscher, Daw, Dayan, & O'Doherty, 2010). The model assumes that participants make choices based on a weighted combination of both model-free and model-based RL algorithms, with a free model-based weight parameter $\omega$ estimated for each participant ($\omega = 1$ reflects a purely model-based agent).

$$V_{s1}^{Net} = \omega \cdot V_{s1}^{MB} + (1-\omega) \cdot V_{s1}^{MF} \qquad [1]$$

The model-free algorithm, based on the temporal difference learning (Rummery & Niranjan, 1994), updates the model-free value of each first-stage option, $V_{s1}^{MF}$, by the reward prediction error (RPE) multiplied by a free first-stage learning rate parameter $\alpha_1$ at the onset of the second stage and reward outcome.

$$V_{s1}^{MF}(t+1) = V_{s1}^{MF}(t) + \alpha_1 \cdot (V_{s2,chosen}(t) - V_{s1}^{MF}(t)) \text{ at the second-stage onset,}$$

$$V_{s1}^{MF}(t+1) = V_{s1}^{MF}(t) + \alpha_1 \cdot (reward - V_{s2}(t)) \text{ at the reward onset.}$$

In contrast, the model-based algorithm computes the utility of each first-stage option by considering the transition structure.

$$V_{s1}^{MB} = p \cdot max(V_3, V_4) + (1-p) \cdot max(V_5, V_6),$$

where $p = 0.7$ for the common transition and $p = 0.3$ for the rare transition. The second-stage options' value is updated only by the model-free algorithm since there is only a reward outcome without further transition in the task. Thus, the model-based and model-free values are identical for the second-stage options.

$$V_{s2}(t+1) = V_{s2}(t) + \alpha_2 \cdot (reward - V_{s2}(t))$$

The model then submits the model-based and model-free value to the softmax function, in which free inverse temperature parameters of each stage (i.e., $\beta_1$, $\beta_2$) determine the stochasticity in the participant's choice, to estimate the probability of choosing each option. For instance, the probability of choosing option 2 in the first stage is computed as:

$$P_2^{stage1} = \frac{1}{1 + exp(-\beta_1 \cdot (V_2^{Net} - V_1^{Net}) - \pi \cdot (C_1 - C_2))}$$

where $V_2^{Net}$ and $V_1^{Net}$ indicate the net value of each option in the first stage, computed by Equation 1 above, and $C_i = 1$ if the previous choice is $i$, otherwise $C_i = 0$. $\pi$ is a free perseverance parameter that captures whether the participant chose the same option as on the previous trial. Thus, a higher $\beta$ value indicates that choices are strongly influenced by the difference in values, and a higher $\pi$ value indicates the participant has a tendency to repeat the same choices. Overall, the model consists of 6 parameters ($\alpha_1, \alpha_{2,} \beta_1, \beta_2, \pi, \omega$).

We separately fitted the model for each group using hierarchical Bayesian modeling, which incorporates group-level information about parameter values to constrain parameter estimation at an individual level. This modeling framework improves the accuracy of parameter estimation (Ahn, Krawitz, Kim, Busemeyer, & Brown, 2011; Lehmann & Casella, 2006; Rouder & Haaf, 2019) by regularizing within-individual variability in parameter estimates (i.e., shrinkage; Efron & Morris, 1977). We then extracted the group-level estimates of the six model parameters and examined group differences for each pair (IGD vs. AUD; IGD vs. HC; AUD vs. HC) (**Figure S3**). To determine that parameter estimates of two groups show a significant difference, 95% highest density interval (HDI) of the subtracted values of each parameter estimate (e.g., $\omega_{IGD} - \omega_{AUD}$) should not include zero.

*Model-based fMRI analysis*

In the first-level analysis, time series of standard RPE estimates were extracted as a model-free regressor. For a model-based regressor, we computed difference regressor representing the residual prediction error not accounted for by model-free RPE values (Daw et al., 2011). The difference regressor was computed by subtracting the model-free RPE values from the hypothetical RPE values that would have occurred if the participant solely used a model-based strategy during the task (Daw et al., 2011). Both the model-free and model-based regressors were included as parametric modulators in the design matrix at the onset of second stage and feedback stage. Additional regressors include timepoints of choice response in the first and second stage, as well as the onset of fixation and first stage. Six motion regressors estimated by fMRIPrep were also added as nuisance regressors to account for head movement.

In the second-level analysis, first-level contrast images were entered into random effects analysis to generate second-level contrasts. Depression and anxiety scores were included as covariates to control for the effect of psychiatric symptoms. The results were thresholded at $p < 0.001$ (uncorrected for multiple comparisons) with an extent threshold of $k \geq 10$ voxels This criterion was adopted to discern potentially meaningful neural activations between the groups, given the exploratory nature of the investigation.

*Functional connectivity analysis - Psychophysiological interaction analysis*

Brain regions related to the salience network were defined based on the Neurosynth (http://neurosynth.org) on 06/22/2023 using terms "salience", "salience network", and "reward' was used to identify relevant brain regions (using a total of 1,444 studies). In the first-level analysis, while using the model-free and model-based RPEs as parametric modulators, we included the 10 nuisance regressors included in the first-level model-based fMRI analysis. In the second-level analysis, first-level contrast images were entered into random effects analysis to generate second-level contrasts. The results were thresholded at $p < 0.001$ (uncorrected) with an extent threshold of $k \geq 10$ voxels and were small-volume-corrected using the brain mask of the salience network.

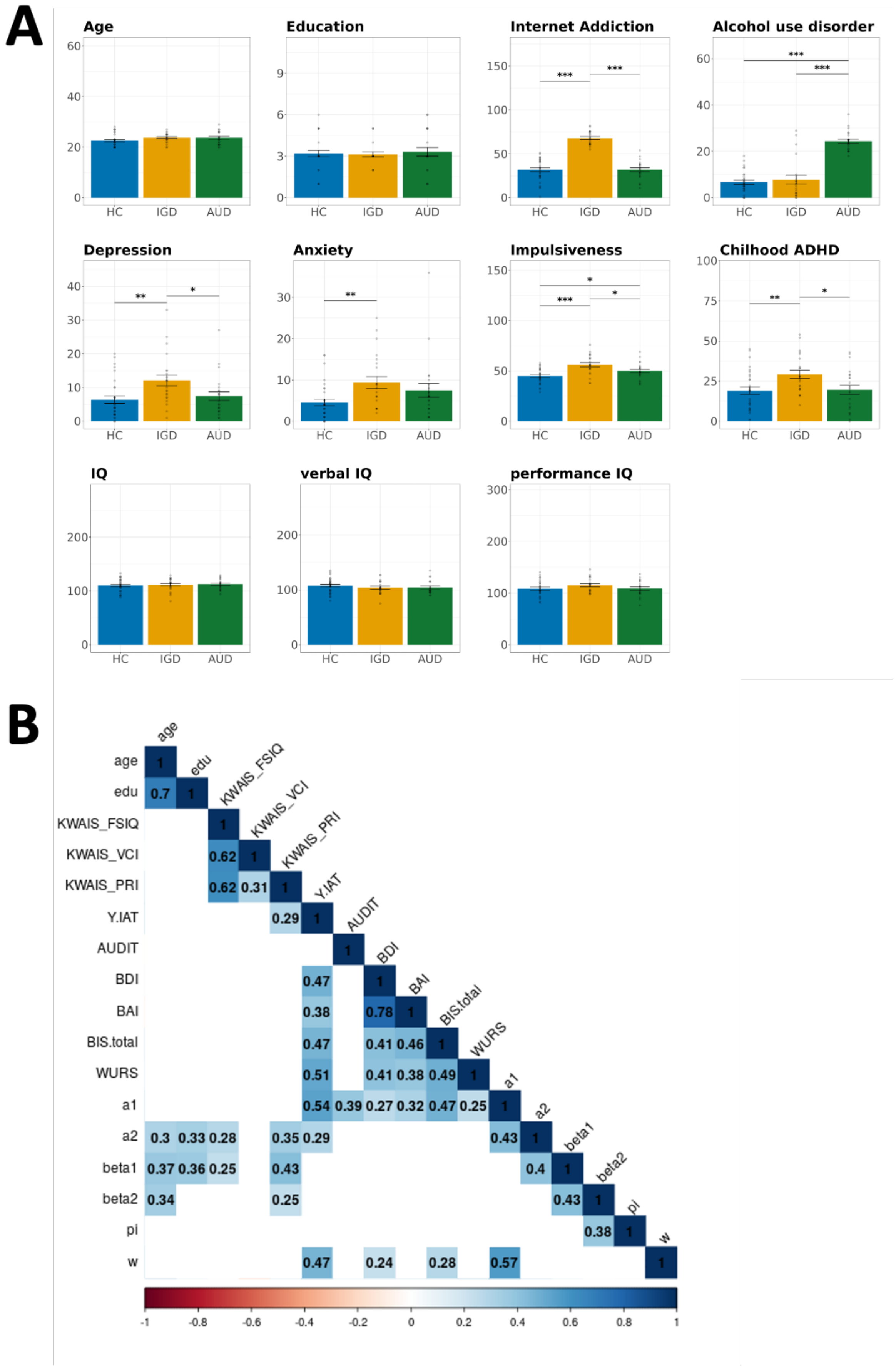
A
Age
Education
Internet Addiction
Alcohol use disorder
Depression
Anxiety
Impulsiveness
Chilhood ADHD
IQ
verbal IQ
performance IQ
HC
IGD
AUD
B
age
edu
KWAIS_FSIQ
KWAIS_VCI
KWAIS_PRI
Y.IAT
AUDIT
BDI
BAI
BIS.total
WURS
a1
a2
beta1
beta2
pi
w

**Figure S1. Group differences and correlations between psychometric measures**

(A) Group differences in psychometric measures. Each bar represents the means of each group, with error bars indicating the 95% confidence interval. Dots correspond to individual participant data. Asterisks denote significant group differences (*, $p < 0.05$; **, $p < 0.005$; ***, $p < 0.001$). Please refer to **Table S1** for detailed values of the measures for each group and the group comparison results. (B) Correlations between the psychometric measures. Each number represents the Pearson's correlation coefficient between the measures corresponding to the x and y axes of each block. Blocks with coefficients indicate a significant correlation between the measures ($p < 0.05$), while blocks without number indicate a nonsignificant correlation.

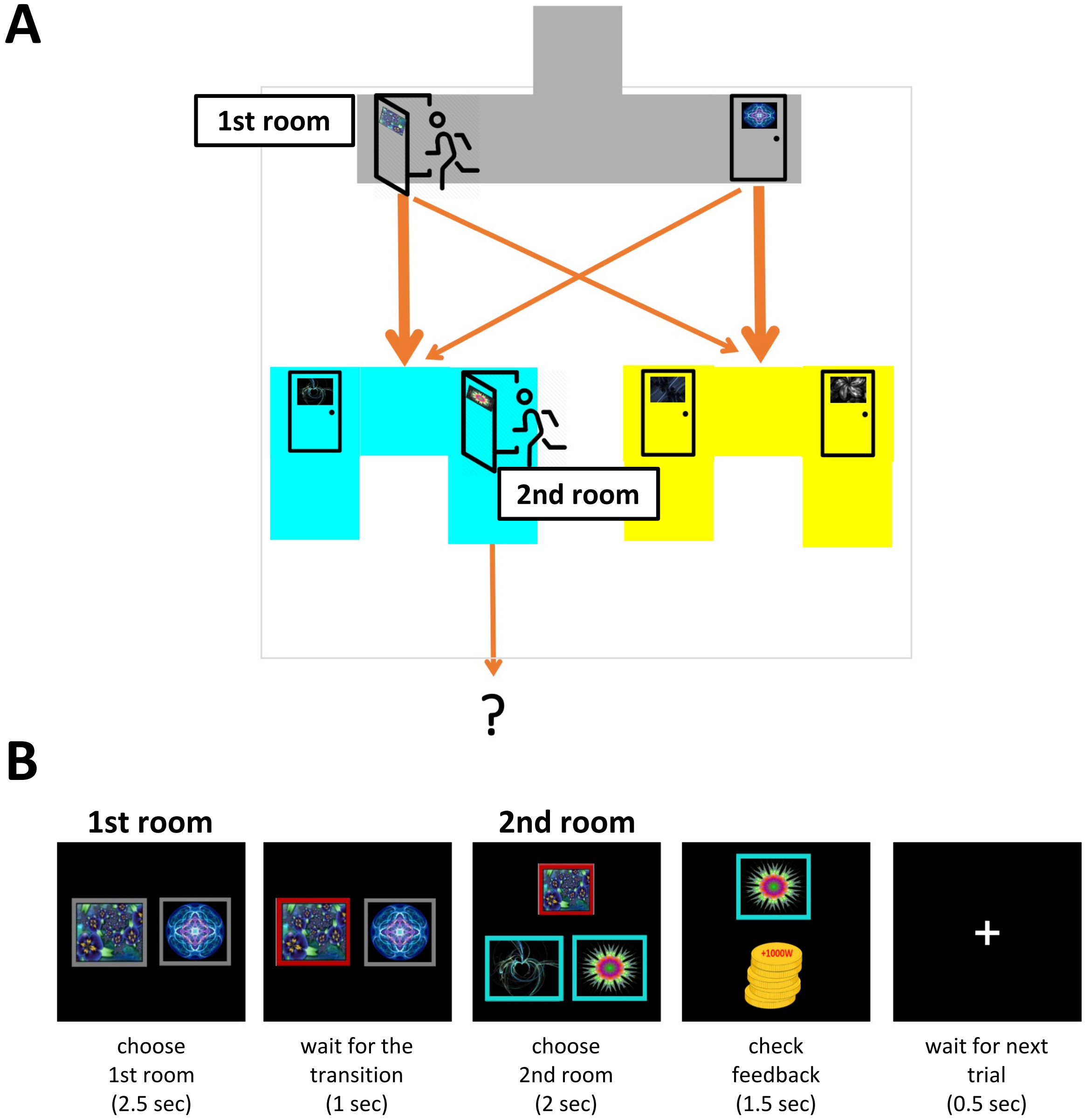


**Figure S2. Schematic figures of the two-stage task used in task instruction**

(A) A schematic figure of exemplar choices used in the task instruction, in which human icon in each stage indicates choices. (B) Process of a single task trial with time points. Inside the fMRI scanner, participants completed 201 task trials, divided into three runs (7.5 seconds per trial, 8.38 minutes per run).

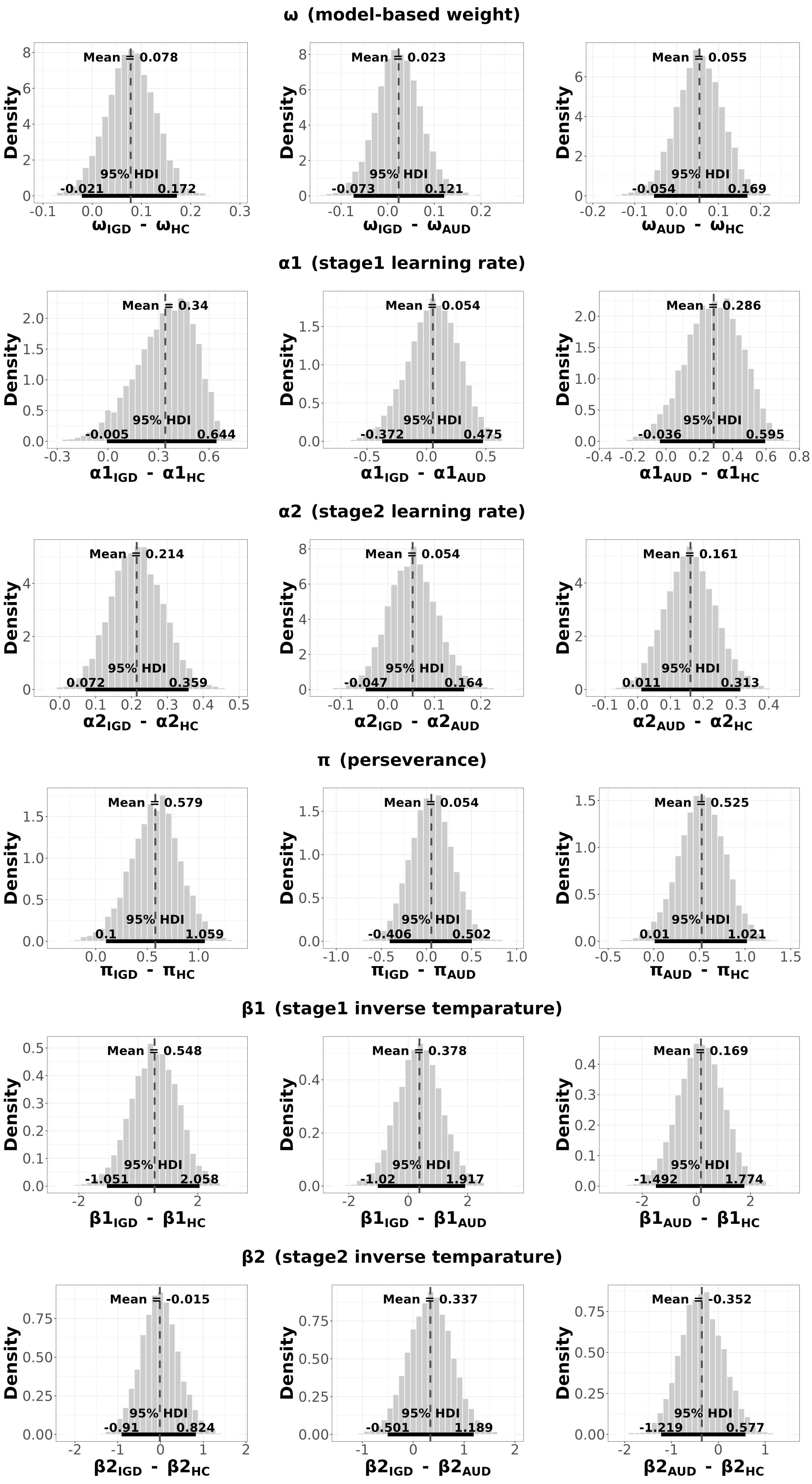
ω (model-based weight)
Mean = 0.078
95% HDI
-0.021
0.172
ωIGD - ωHC
Mean = 0.023
95% HDI
-0.073
0.121
ωIGD - ωAUD
Mean = 0.055
95% HDI
-0.054
0.169
ωAUD - ωHC
α1 (stage1 learning rate)
Mean = 0.34
95% HDI
-0.005
0.644
α1IGD - α1HC
Mean = 0.054
95% HDI
-0.372
0.475
α1IGD - α1AUD
Mean = 0.286
95% HDI
-0.036
0.595
α1AUD - α1HC
α2 (stage2 learning rate)
Mean = 0.214
95% HDI
0.072
0.359
α2IGD - α2HC
Mean = 0.054
95% HDI
-0.047
0.164
α2IGD - α2AUD
Mean = 0.161
95% HDI
0.011
0.313
α2AUD - α2HC
π (perseverance)
Mean = 0.579
95% HDI
0.1
1.059
πIGD - πHC
Mean = 0.054
95% HDI
-0.406
0.502
πIGD - πAUD
Mean = 0.525
95% HDI
0.01
1.021
πAUD - πHC
β1 (stage1 inverse temparature)
Mean = 0.548
95% HDI
-1.051
2.058
β1IGD - β1HC
Mean = 0.378
95% HDI
-1.02
1.917
β1IGD - β1AUD
Mean = 0.169
95% HDI
-1.492
1.774
β1AUD - β1HC
β2 (stage2 inverse temparature)
Mean = -0.015
95% HDI
-0.91
0.824
β2IGD - β2HC
Mean = 0.337
95% HDI
-0.501
1.189
β2IGD - β2AUD
Mean = -0.352
95% HDI
-1.219
0.577
β2AUD - β2HC
Density

**Figure S3. Distributions of group differences in model parameter estimates**

To assess group differences in the model parameter estimates, we computed the differences between each pair (HC vs. IGD; IGD vs. AUD; HC vs. AUD). The distributions represent the subtracted values of each parameter estimate from one group to the other group (e.g., $\omega_{IGD} - \omega_{AUD}$). The black lines under the distributions indicate the 95% highest density interval (HDI). Group differences were considered significant when the 95% HDI did not include zero.

### Table S1. Group differences in psychometric measures

| | HC | IGD | AUD | F (*p* value) |
|---|---|---|---|---|
| **Sample size** | N = 30 | N = 22 | N = 22 | |
| **Age** | 22.6 (22.14, 23.06) | 23.73 (23.33, 24.13) | 23.73 (23.18, 24.28) | 2.032 (p = 0.139) |
| **Education** | 3.2 (2.98, 3.42) | 3.14 (2.95, 3.33) | 3.32 (3, 3.64) | 0.126 (p = 0.882) |
| **Internet addiction (IAT)** | 32.03 (29.67, 34.39) | 68 (66.34, 69.66) | 32.05 (29.83, 34.26) | 84.716 (p < 0.001) |
| **Alcohol use (AUDIT)** | 6.7 (5.86, 7.54) | 7.82 (5.96, 9.68) | 24.36 (23.44, 25.29) | 62.578 (p < 0.001) |
| **Depression (BDI)** | 6.37 (5.3, 7.43) | 12.14 (10.52, 13.75) | 7.45 (6.14, 8.77) | 5.358 (p < 0.05) |
| **Anxiety (BAI)** | 4.57 (3.73, 5.4) | 9.41 (7.98, 10.84) | 7.45 (5.81, 9.1) | 3.906 (p < 0.05) |
| **Impulsivity (BIS)** | 44.93 (43.57, 46.29) | 56.18 (54.06, 58.3) | 50.05 (48.3, 51.79) | 11.205 (p < 0.001) |
| **Childhood ADHD (WURS)** | 18.97 (16.71, 21.22) | 29.23 (26.62, 31.83) | 19.55 (16.73, 22.36) | 4.914 (p < 0.05) |
| **IQ (K-WAIS)** | 110.43 (108.24, 112.63) | 111.64 (109.07, 114.21) | 112.45 (110.36, 114.55) | 0.206 (p = 0.814) |
| **verbal IQ (VCI)** | 107.47 (104.97, 109.97) | 103.86 (101.27, 106.45) | 104.09 (101.52, 106.66) | 0.667 (p = 0.516) |
| **performance IQ (PRI)** | 108.73 (106, 111.47) | 115.27 (112.38, 118.17) | 109.05 (105.87, 112.22) | 1.501 (p = 0.23) |

*Note.* The data presented in the first three columns represent the means (95% confidence interval) for each group (HC, IGD, and AUD), or the sample size. The results of a one-way analysis of variance (ANOVA) with group factors (i.e., HC, IGD, AUD) are shown in the rightmost column, including the F-statistics and p value. The abbreviations used in the table are as follows: IAT, Young Internet Addiction Test; AUDIT, Alcohol Use Disorder Identification Test; BDI, Beck Depression Inventory; BAI, Beck Anxiety Inventory; BIS, Barratt Impulsiveness Scale; WURS, Wender Utah Rating Scale; K-WAIS, Korean version of the Wechsler Adult Intelligence Scale; VCI, Verbal Comprehension Index Scale; PRI, Perceptual Reasoning Index Scale.

**Table S2. Model comparison results**

| | HC | IGD | AUD |
|---|---|---|---|
| **4 parameter model** | 11907.628 | 7779.664 | 8104.869 |
| **6 parameter model** | **11838.170** | **7711.565** | **8062.970** |
| **7 parameter model** | 11840.879 | 7715.729 | 8064.469 |

*Note.* The table presents the leave-one-out information criterion (LOOIC) values for three different models: a seven-parameter model, a six-parameter model, and a four-parameter model, for each group (HC, healthy control; IGD, internet gaming disorder; AUD, alcohol use disorder). The six-parameter model, highlighted in blue, exhibited the lowest LOOIC value, indicating the best model fit among the three models.

### Table S3. Results of linear mixed effects logistic regression for stay probability

| | HC | | IGD | | AUD | |
|---|---|---|---|---|---|---|
| | ***Coef***<br>**[95% CI]** | ***p* value** | ***Coef***<br>**[95% CI]** | ***p* value** | ***Coef***<br>**[95% CI]** | ***p* value** |
| **Reward** | -0.778<br>[-1.129, -0.427] | *p*=1.40e-05 | -1.117<br>[-1.440, -0.794] | *p*=1.19e-11 | -1.119<br>[-1.536, -0.702] | *p*=1.44e-07 |
| **Transition probability** | -1.077<br>[-1.406, -0.748] | *p*=1.40e-10 | -1.055<br>[-1.374, -0.735] | *p*=9.38e-11 | -1.238<br>[-1.954, -0.523] | *p*=0.001 |
| **Reward x transition probability** | 1.970<br>[1.661, 2.279] | *p*<2.00e-16 | 3.311<br>[2.912, 3.709] | *p*<2.00e-16 | 3.204<br>[-2.776, 3.631] | *p*<2.00e-16 |

*Note.* The table presents the coefficients, 95% confidence intervals (CI), and p-values for the main effects of reward, transition probability, and the interaction between the two. The residual degree of freedom of each group are as follows: HC = 5,969, IGD = 4,370, AUD = 4,385.

### Table S4. Second-level results of the model-based fMRI

| Group | Corresponding Brain Region | Peak MNI Coordinates (x, y, z) | T value | Z value | *p* value (peak voxel) | *p* value (cluster level) | cluster size |
|---|---|---|---|---|---|---|---|
| **HC** | Fusiform gyrus (L) | -36, -84, -8 | 4.82 | 4.01 | $p < 0.001^{*}$ | $p = 0.102$ | 10 |
| **IGD** | Insula (R) | 42, 14, -14 | 4.56 | 3.64 | $p < 0.001^{*}$ | $p = 0.085$ | 15 |
| **AUD** | Orbitofrontal cortex (R) | 35, 55, -8 | 6.39 | 4.56 | $p < 0.001^{*}$ | $p = 0.016^{\dagger}$ | 29 |
| | Occipital lobe (R) | 31, -91, -4 | 5.15 | 3.98 | $p < 0.001^{*}$ | $p = 0.088$ | 13 |
| | Posterior cingulate cortex | 1, -35, 36 | 5.05 | 3.93 | $p < 0.001^{*}$ | $p = 0.013^{\dagger}$ | 31 |
| | Occipital lobe (L) | -22, -99, -4 | 4.98 | 3.90 | $p < 0.001^{*}$ | $p = 0.088$ | 13 |
| | Anterior cingulate cortex | -3, 44, 6 | 4.91 | 3.86 | $p < 0.001^{*}$ | $p = 0.019^{\dagger}$ | 27 |

*Note.* The table displays significant results of the second-level one-sample t-tests conducted for the model-based fMRI analysis, using a model-based reward prediction error (RPE) as a parametric modulator ($p < 0.001$, uncorrected; cluster size, $k \geq 10$). The asterisks (*) indicate significant results with $p < 0.001$, and the cross symbols (†) indicate significant results with $p < 0.05$.

**Table S5. PPI second-level results of the internet gaming disorder (IGD) group**

| Seed region | Corresponding Brain Region | Peak MNI Coordinates (x, y, z) | T value | Z value | *p* value (uncorrected) | *p* value (cluster level) | cluster size |
|---|---|---|---|---|---|---|---|
| Insula (R) | Putamen (R) | 31, 2, -14 | 5.23 | 3.98 | $p < 0.001^{*}$ | $p = 0.014^{\dagger}$ | 24 |
| | Insula (L) | -48, 10, -8 | 5.75 | 4.23 | $p < 0.001^{*}$ | $p = 0.346$ | 3 |
| | Occipital lobe | -48, -76, -4 | 4.37 | 3.53 | $p < 0.001^{*}$ | $p = 0.601$ | 1 |
| | | 46, -72, -4 | 4.31 | 3.50 | $p < 0.001^{*}$ | $p = 0.110$ | 9 |
| | | -48, -69, 6 | 4.23 | 3.45 | $p < 0.001^{*}$ | $p = 0.445$ | 2 |
| Insula (L) | Anterior Cingulate Cortex | 5, 10, 32 | 4.24 | 3.45 | $p < 0.001^{*}$ | $p = 0.012^{\dagger}$ | 24 |
| | Superior temporal gyrus (R) | 31, -9, -4 | 3.67 | 3.11 | $p = 0.001^{\dagger}$ | $p = 0.587$ | 1 |
| | Occipital lobe | 38, -50, -18 | 5.14 | 3.94 | $p < 0.001^{*}$ | $p = 0.119$ | 8 |
| | | -40, -69, -14 | 4.82 | 3.78 | $p < 0.001^{*}$ | $p = 0.330$ | 3 |
| | | -36, -88, -4 | 3.92 | 3.27 | $p = 0.001^{\dagger}$ | $p = 0.429$ | 2 |
| | | -40, -58, -14 | 3.89 | 3.24 | $p = 0.001^{\dagger}$ | $p = 0.429$ | 2 |
| | | 8, -84, -4 | 3.82 | 3.2 | $p = 0.001^{\dagger}$ | $p = 0.429$ | 2 |
| | | -40, -80, -8 | 3.72 | 3.14 | $p = 0.001^{\dagger}$ | $p = 0.587$ | 1 |

*Note.* The table presents the significant psychophysiological interaction (PPI) analysis results obtained from the second-level one-sample t-tests conducted for the IGD group, using a model-based reward prediction error (RPE) as the psychological variable ($p < 0.001$, uncorrected; cluster size, $k \geq 10$). After applying small volume correction using an extent brain mask created by combining meta-analysis brain maps of "salience", "salience network", and "reward" extracted from the Neurosynth, the resulting clusters within the specified volume displayed smaller sizes than the initially applied threshold of $k \geq 10$. The asterisks (*) indicate results with $p < 0.001$, and the cross symbols (†) indicate results with $p < 0.05$.

#### Table S6. Second-level group comparison results of the PPI analysis

| Group | Seed region | Corresponding Brain Region | Peak MNI Coordinates (x, y, z) | T value | Z value | *p* value (uncorrected) | *p* value (cluster level) | cluster size |
|---|---|---|---|---|---|---|---|---|
| Insula (R) | HC < IGD | Insula (L) | -48, 10, -8 | 3.89 | 3.59 | $p < 0.001^{*}$ | $p = 0.513$ | 2 |
| | | Parietal frontal lobe (R) | 35, 6, 56 | 3.61 | 3.36 | $p < 0.001^{*}$ | $p = 0.292$ | 5 |
| | | Premotor cortex (R) | 27, -5, 52 | 3.31 | 3.11 | $p < 0.001^{*}$ | $p = 0.657$ | 1 |
| | | Fusiform gyrus (R) | 46, -54, -18 | 3.75 | 3.47 | $p < 0.001^{*}$ | $p = 0.657$ | 1 |
| | | | 31, -61, -18 | 3.34 | 3.14 | $p < 0.001^{*}$ | $p = 0.657$ | 1 |
| | | Parietal lobe (R) | 46, -31, 42 | 3.66 | 3.4 | $p < 0.001^{*}$ | $p = 0.657$ | 1 |
| | | Occipital lobe | 31, -84, -8 | 3.67 | 3.41 | $p < 0.001^{*}$ | $p = 0.347$ | 4 |
| | | | 5, -69, 46 | 3.55 | 3.31 | $p < 0.001^{*}$ | $p = 0.513$ | 2 |
| | | | -40, -58, -18 | 3.54 | 3.31 | $p < 0.001^{*}$ | $p = 0.417$ | 3 |
| | | | -14, -72, 52 | 3.54 | 3.31 | $p < 0.001^{*}$ | $p = 0.657$ | 1 |
| | | | 12, -84, 6 | 3.51 | 3.28 | $p < 0.001^{*}$ | $p = 0.513$ | 2 |
| | | | -6, -84, -4 | 3.44 | 3.22 | $p < 0.001^{*}$ | $p = 0.513$ | 2 |
| | AUD < IGD | Occipital lobe | -10, -95, -4 | 4.41 | 3.92 | $p < 0.001^{*}$ | $p = 0.083$ | 13 |
| | | | 12, -84, 2 | 4.09 | 3.69 | $p < 0.001^{*}$ | $p = 0.392$ | 3 |
| | | | -25, -95, 6 | 3.57 | 3.29 | $p < 0.001^{*}$ | $p = 0.637$ | 1 |
| | | | 31, -95, -4 | 3.52 | 3.25 | $p < 0.001^{*}$ | $p = 0.321$ | 4 |
| Insula (L) | AUD < IGD | Occipital lobe | 12, -95, 12 | 3.79 | 3.46 | $p < 0.001^{*}$ | $p = 0.492$ | 2 |
| | | | -25, -95, 6 | 3.41 | 3.16 | $p < 0.001^{*}$ | $p = 0.639$ | 1 |
| | AUD > IGD | Posterior cingulate cortex | -3, -46, 32 | 3.81 | 3.47 | $p < 0.001^{*}$ | $p = 0.271$ | 5 |

*Note.* The table presents the results of the second-level group comparisons using two-sample t-tests in the psychophysiological interaction (PPI) analysis ($p < 0.001$, uncorrected; cluster size, $k \geq 10$). After applying small volume correction using an extent brain mask created by combining meta-analysis brain maps of "salience", "salience network", and "reward" extracted from the Neurosynth, the resulting clusters within the specified volume displayed smaller sizes than the initially applied threshold of $k \geq 10$. The asterisks (*) indicate results with $p < 0.001$, and the cross symbols (†) indicate results with $p < 0.05$.
